# Sex-specific vasopressin signaling buffers stress-dependent synaptic changes in female mice

**DOI:** 10.1101/2020.04.30.070532

**Authors:** Spencer P. Loewen, Dinara Baimoukhametova, Jaideep S. Bains

## Abstract

In many species, social networks provide benefit for both the individual and the collective. In addition to transmitting information to others, social networks provide an emotional buffer for distressed individuals. Our understanding about the cellular mechanisms that contribute to buffering is poor. Stress has consequences for the entire organism, including a robust change in synaptic plasticity at glutamate synapses onto corticotropin-releasing hormone (CRH) neurons in the paraventricular nucleus of the hypothalamus (PVN). In females, however, this stress-induced metaplasticity is buffered by the presence of a naïve partner. This buffering may be due to discrete behavioral interactions, signals in the context in which the interaction occurs (i.e. olfactory cues), or it may be influenced by local signaling events in the PVN. Here, we show that local vasopressin (VP) signaling in PVN buffers the short-term potentiation (STP) at glutamate synapses after stress. This social buffering of metaplasticity, which requires the presence of another individual, was prevented by pharmacological inhibition of the VP 1a receptor in female mice. Exogenous VP mimicked the effects of social buffering and reduced STP in CRH^PVN^ neurons from females but not males. These findings implicate VP as a potential mediator of social buffering in female mice.

**Significance Statement:** In many organisms, including rodents and humans, social groups are beneficial to overall health and well-being. Moreover, it is through these social interactions that the harmful effects of stress can be mitigated—a phenomenon known as social buffering. In the present study, we describe a critical role for the neuropeptide vasopressin in social buffering of synaptic metaplasticity in stress-responsive corticotropin-releasing hormone neurons in female mice. These effects of vasopressin do not extend to social buffering of stress behaviors, suggesting this is a very precise and local form of sex-specific neuropeptide signaling.

## Introduction

Many species, including rodents and humans, transmit and receive information through social interactions. These interactions are vital for many functions, including survival, reproduction, resource allocation, and affiliation (Krause et al., 2002; Silk, 2007). This complexity is one reason our understanding of this critical biological processes is limited. Understanding social behavior, and, specifically, social behavior in specific circumstances, requires a detailed understanding of the brain at multiple levels from circuits to synapses to cells. In many species, social behaviors show sexual divergence; males and females show distinct features of social behavior related to parenting (Lonstein and De Vries, 2000), aggression (Archer, 2009), affiliative behavior (Taylor et al., 2000), and empathy (Christov-Moore et al., 2014). The neuropeptides oxytocin (OT) and vasopressin (VP) play critical roles in many social behaviors (Johnson and Young, 2017), and there has been increasing attention given to the role of OT and VP as sexually dimorphic social signals (de Vries, 2008; Dumais and Veenema, 2016). How they specifically modulate neuronal or synaptic function in a sex-specific fashion within the context of the social brain, however, is not clear.

Social interactions also leave enduring marks on the brain. We have recently shown that social interactions can transmit negative affective states and cause changes at the level of glutamate synapses in first and second order recipients of stress (Sterley et al., 2018). These changes result in a stress-induced metaplasticity at glutamate synapses onto corticotropin-releasing hormone (CRH) neurons in the hypothalamic paraventricular nucleus (PVN) (Sterley et al., 2018), which are adjacent to both OT and VP neurons within this structure (Biag et al., 2012). Intriguingly, social interactions also result in a decrease in the synaptic changes in female individuals subjected directly to stress (Sterley et al., 2018). Conversely, both social isolation and acute stress have similar effects on the biophysical properties of CRH^PVN^ neurons in females (Senst et al., 2016). These findings suggest that the long-term consequences of stress are impacted by social interactions in a sexually dimorphic fashion. However, the mechanisms responsible for the sex-specific curtailment of synaptic changes following social interaction are not known.

Multiple lines of evidence suggest that neurons in the PVN are sensitive to neuromodulation by locally released substances (Landgraf and Neumann, 2004; Ludwig and Leng, 2006). These substances, which include CRH (Kuzmiski et al., 2010; Ramot et al., 2017; Jiang et al., 2018), dynorphin (Iremonger and Bains, 2009), enkephalin (Wamsteeker Cusulin et al., 2013a), OT (Kombian et al., 1997; Oliet et al., 2007), and VP (Ludwig, 1998), are released from the somatodendritic compartments of PVN neurons. Intranuclear release of VP in the PVN following social defeat or forced swimming exerts an inhibitory effect on hypothalamic-pituitary-adrenal (HPA) axis activation and subsequent ACTH secretion (Wotjak et al., 1996, 1998). Dendritically released VP from magnocellular neurosecretory cells can affect the excitability of preautonomic neurons up to 100 μm away (Son et al., 2013). These cellular effects, combined with the involvement of VP in pair-bonding and affiliative behaviors (Young and Wang, 2004; Insel, 2010), the demonstration that VP neurons in the PVN are selectively targeted by a synaptic relay from olfactory bulb neurons that express a family of olfactory receptors (OR37) that buffer the effects of stress (Bader et al., 2012; Klein et al., 2015), and the adjacent cytoarchitectural arrangement of CRH and VP neurons in the PVN (Biag et al., 2012), led us to hypothesize that local release of VP in the PVN may contribute to social buffering of synaptic metaplasticity in females.

Here, we demonstrate that the presence of others reduces stress-induced grooming behavior and metaplasticity at glutamate synapses onto CRH^PVN^ neurons in female mice. The effects on synapses, but not behavior, require local actions of VP at VP 1a receptors (V1aRs) on CRH^PVN^ neurons, suggesting that VP has sex-specific neuromodulatory effects that ‘erase’ the synaptic effects of a stressful experience in females.

## Materials and Methods

### Animals

All animal protocols conformed to the standards of the Canadian Council on Animal Care and were approved by the University of Calgary Animal Care and Use Committee. Experiments were performed on 3- to 5-week old male and female *Crh-IRES-Cre; Ai14* mice in which CRH neurons express a tdTomato fluorophore (Wamsteeker Cusulin et al., 2013b). Mice were housed in whole litters on a 12-h:12-h light:dark cycle (lights on at 6:00 a.m.) and were provided with food and water *ad libitum.* At least sixteen hours prior to acute stress or slice preparation, mice were housed either alone or in same-sex littermate pairs, depending on the experiment. Mice were randomly assigned to the experimental groups. Stress was induced between 9:30 and 10:30 a.m. during the light phase by subjecting mice to a footshock (FS) protocol consisting of a 0.5 mA FS for 2 s delivered every 30 s over a 5-min period. In experiments involving injection of the selective V1aR antagonist, SR 49059 (SR, Sigma-Aldrich), SR was dissolved in polyethylene glycol (PEG) and injected i.p. 5 min prior to FS at a dose of 2 mg/kg in a final volume of 50 μl. Control mice received a 50 μl vehicle injection of PEG. Mice were video-recorded for 30 min following return of the FS mouse to the homecage. Videos were analyzed and scored for grooming, anogenital sniffing (directing the snout toward the anogenital region of the conspecific), and head/torso sniffing (directing the snout toward the head or torso of the conspecific). Social discrimination indices were generated by dividing the amount of time spent sniffing the anogenital region of the conspecific by the total amount of time spent sniffing both the anogenital and head/torso regions of the conspecific.

### Slice preparation

Immediately following FS or the 30-min social interaction, mice were anesthetized with isofluorane and decapitated. Brains were rapidly removed and immersed in ice-cold (0–4°C) slicing solution consisting of the following (in mM): 87 NaCl, 2.5 KCl, 0.5 CaCl_2_, 7 MgCl_2_, 25 NaHCO_3_, 25 D-glucose, 1.25 NaH_2_PO_4_, 75 sucrose, and saturated with 95% O_2_/5% CO_2_. Brains were blocked and 250-μm coronal sections containing the PVN were obtained using a vibratome (VT1200S; Leica). Slices recovered for 1 h in 30°C aCSF consisting of the following (in mM): 126 NaCl, 2.5 KCl, 26 NaHCO_3_, 2.5 CaCl_2_, 1.5 MgCl_2_, 1.25 NaH_2_PO_4_, 10 glucose, and saturated with 95% O_2_/5% CO_2_. Following the 1-hr recovery period, slices treated with VP were transferred to a separate chamber containing 100 nM VP dissolved in aCSF for 30 min prior to recording. In a subset of experiments, slices were pre-incubated in 10 μM SR dissolved in aCSF for 10 min prior to incubation in both VP and SR for an additional 30 min.

### Electrophysiology

Slices were transferred to a recording chamber superfused with 30–32°C aCSF at a flow rate of 1–2 ml/min. All recordings took place in aCSF containing 100 μM picrotoxin unless otherwise stated. Neurons were visualized using a 40× water-immersion objective mounted on an upright microscope (BX51WI, Olympus) fitted with differential interference contrast optics, a metal halide fluorescence lamp (X-Cite Series 120, EXFO), and a digital camera (OLY-150, Olympus). CRH^PVN^ neurons were identified based on their location and tdTomato fluorescence. Borosilicate glass micropipettes were pulled on a horizontal micropipette puller (P-97, Sutter Instruments) to tip resistances between 2.5–5 MΩ when filled with the intracellular solution consisting of the following (in mM): 108 K-gluconate, 2 MgCl_2_, 8 Na-gluconate, 8 KCl, 1 K_2_-EGTA, 4 K_2_-ATP, and 0.3 Na_3_-GTP buffered with 10 HEPES. EPSCs were evoked 50 ms apart at 0.2-Hz intervals using a monopolar aCSF-filled electrode placed in the neuropil surrounding the cell (~20μM). High-frequency stimulation (HFS) consisted of four 100-Hz stimulations for 1 s every 10 s. AMPAR-mediated EPSCs were isolated by holding the postsynaptic neuron at −70 mV to block voltage-dependent postsynaptic NMDARs. NMDAR-mediated currents were recorded at a holding potential of +40 mV using an intracellular solution consisting of the following (in mM): 130 CsCl, 10 NaCl, 10 EGTA, 0.1 CaCl_2_, 4 K_2_-ATP and 0.3 Na_3_-GTP buffered with 10 HEPES. In some experiments, GTP was replaced with 1 mM guanosine 5’-[β-thio]diphosphate trilithium salt (GDPβS, Sigma-Aldrich). In the indicated experiments, VP was applied to slices via syringe pump (R-99, Razel Scientific Instruments) or focally to cells via a puff pipette (20 psi, 1 s) connected to a Picospritzer II microcellular injection unit (General Valve). Access resistance (<20 MΩ) was continuously monitored throughout the experiments; recordings were excluded from analysis if it exceeded a 15% change.

### Data analysis and statistics

Signals were amplified using a MultiClamp 700B amplifier (Molecular Devices), low-pass filtered at 1 kHz, and digitized at 10 kHz using the Digidata 1440A (Molecular Devices). Data were recorded (pClamp 10.4, Molecular Devices) and stored on a computer for offline analysis. Evoked EPSC amplitude was calculated from the baseline current prior to stimulation to the peak synaptic current. The magnitude was calculated as the average amplitude of 12 evoked EPSCs immediately following HFS (0–1min post-HFS). The mean firing frequency and membrane potential values were calculated from the 1 min baseline period immediately prior to VP application and a 20 s period around the peak effect. The AMPA/NMDA ratio was determined by averaging 15 AMPAR-mediated EPSC traces and 15 NMDAR-mediated EPSC traces per cell. Statistical analyses were performed using GraphPad Prism 6.04. Two-tailed *t* tests were used to compare post-HFS to 100% baseline. Unpaired *t* tests were used to compare the means to two independent groups. Paired *t* tests were used to compare the means of two dependent groups. A one-way ANOVA was used to compare the means of multiple groups, followed by Tukey’s or Sidak’s post hoc, multiple comparisons tests. Data are presented as mean ± SEM.

## Results

### Social buffering of stress-induced grooming behavior and STP

CRH^PVN^ neurons play a crucial role in multiple stress-related behaviors (Füzesi et al., 2016; Zhang et al., 2017; Sterley et al., 2018; Daviu et al., 2020). Immediately after an acute stress, CRH^PVN^ neurons, through collateral projections to the lateral hypothalamus, drive self-grooming (Füzesi et al., 2016). We assessed whether the presence of a partner attenuates grooming. A single-housed female mouse (Fig. 1*A*) or one mouse from a female dyad of littermates (Fig. 1*B*) was subjected to FS stress and then returned to its homecage for 30 min; spontaneous behavior of both mice was recorded during this time. Single-housed female mice spent more time grooming than pair-housed female mice (single-housed, 511.8 ± 34.3 s, *N* = 3; pair-housed, 298.4 ± 38.7 s, *N* = 5; *p* = 0.0098, unpaired *t* test, two-tailed; Fig. 1*Ci–E*).

**Figure 1.**
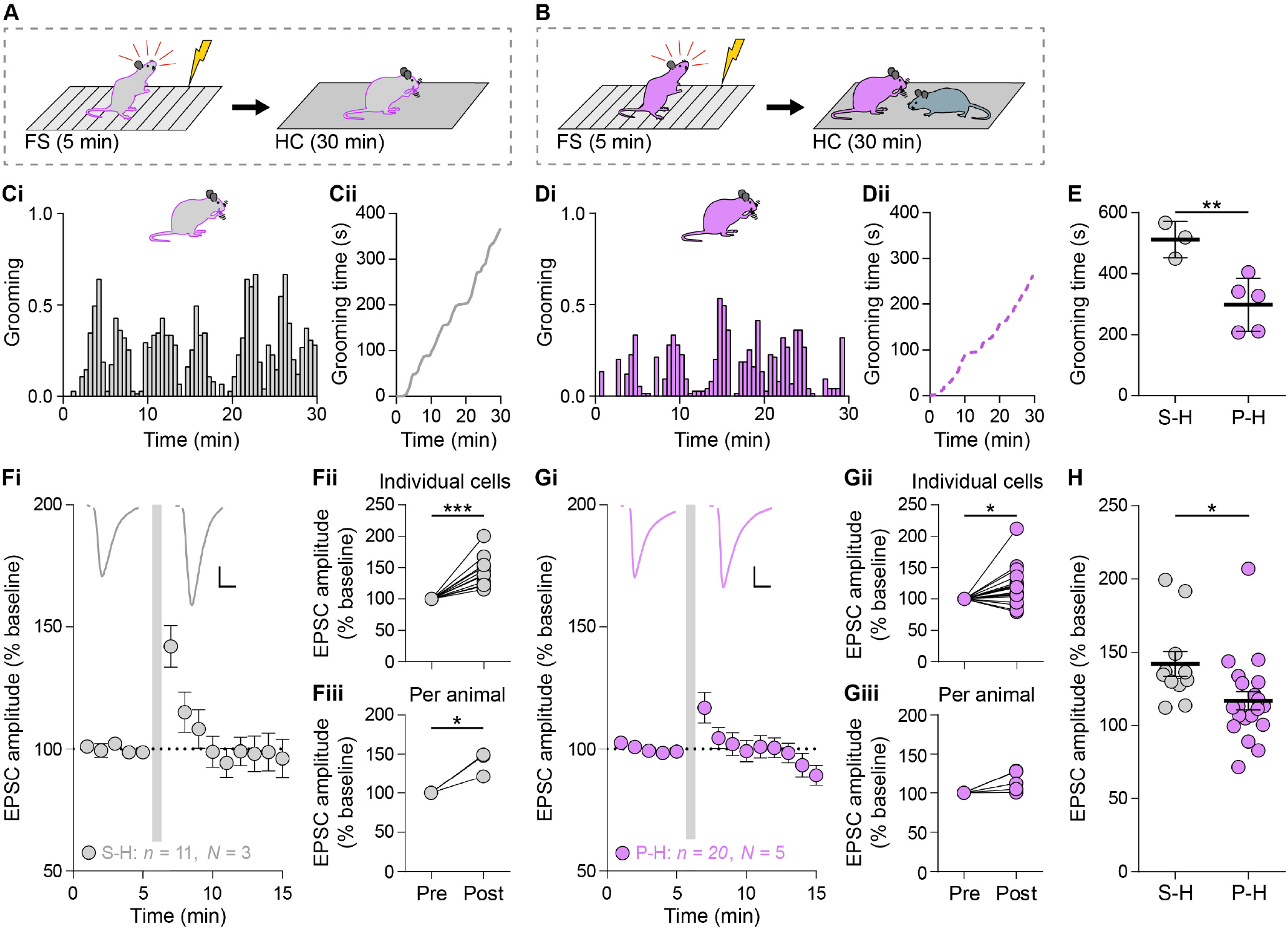
Social interaction reduces stress-induced grooming behavior and STP in CRH^PVN^ neurons. **A)** Single-housed female mice were subjected to the FS protocol (0.5 mA for 2 s every 30 s over 5 min) and then returned to the homecage for 30 min. **B)** One mouse from a female dyad of littermates was exposed to FS and returned to its partner in the homecage for 30 min. **Ci)** Histogram showing the amount of self-grooming displayed by single-housed FS mice (*N* = 3) during the 30-min observation period. **Cii)** Cumulative graph illustrates the relative grooming of single-housed FS mice. **Di)** Histogram showing the amount of self-grooming displayed by pair-housed FS mice (*N* = 5) during the 30-min observation period. **Dii)** Cumulative graph illustrates the relative grooming of pair-housed FS mice. **E)** Single-housed FS mice engaged in more selfgrooming than pair-housed FS mice (single-housed, mean: 511.8 ± 34.3 s, *N* = 3 mice; pair-housed, mean: 298.4 ± 38.7 s, *N* = 5 mice; *p* = 0.0098, unpaired *t* test, two-tailed). Horizontal bars represent the means. **Fi)** EPSCs in CRH^PVN^ neurons of single-housed FS mice potentiated following HFS (gray bar) relative to baseline. Inset, synaptic currents before and after HFS in single-housed FS mice. **Fii)** STP was present in cells from single-housed FS mice (mean: 142.0 ± 8.6%, *n* = 11, *p* = 0.0002 vs. baseline, one-sample *t* test, two-tailed). **Fiii)** STP was present in single-housed FS mice (mean: 138.1 ± 8.7%, *n* = 3, *p* = 0.04 vs. baseline, one-sample *t* test, twotailed). **Gi)** EPSCs in CRH^PVN^ neurons of pair-housed FS mice potentiated following HFS (gray bar) relative to baseline. Inset, synaptic currents before and after HFS in pair-housed FS mice. **Gii)** STP was present in cells from pair-housed FS mice (mean: 116.9 ± 6.3%, *n* = 20, *p* = 0.01 vs. baseline, one-sample *t* test, two-tailed). **Giii)** STP was absent in pair-housed FS mice (mean: 110.8 ± 7.1%, *n* = 5, *p* = 0.06 vs. baseline, one-sample *t* test, two-tailed). **H)** STP (average ESPC amplitude first minute post-HFS relative to baseline, individual cells shown) was larger in single-housed FS mice than in pair-housed FS mice (*p* = 0.02, unpaired *t* test, two-tailed). Horizontal bars represent the means. Scale bars (**Fi,Gi**) represent 5 ms and 20 pA. Inset currents (**Fi,Gi**) before HFS are scaled to allow for easier comparison after HFS. Error bars represent ± SEM.

Next, we conducted experiments in single- and pair-housed female mice to ask whether the presence of a partner decreases the synaptic consequences of stress in female mice. Immediately following the 30-min observation period described above, we prepared brain slices from the animals and performed subsequent whole-cell recordings on CRH^PVN^ neurons to assess STP. Following high-frequency stimulation of the neuropil, glutamate synapses onto CRH^PVN^ neurons from single-housed female mice subjected to FS show robust STP (142.0 ± 8.6%, *n* = 11, *p* = 0.0002 vs. baseline; Fig. 1*Fi,Fii*); STP was also observed in pair-housed female FS mice (116.9 ± 6.3%, *n* = 20, *p* = 0.01 vs. baseline; Fig. 1*Gi,Gii*), but this was significantly reduced when compared to single-housed mice (*p* = 0.02, unpaired *t* test, two-tailed; Fig. 1*H*). Meanwhile, per animal analysis revealed that STP was present in single-housed FS mice (138.1 ± 8.7%, *N* = 3, *p* = 0.04 vs baseline; Fig. 1*Fiii*) but not in pair-housed FS mice (110.8 ± 7.1%, *N* = 5, *p* = 0.06 vs. baseline; Fig. 1*Giii*). These findings are consistent with previous findings from our laboratory (Sterley et al., 2018), and provide a useful model to study the neural consequences of social buffering.

### VP receptor signaling is required for social buffering of stress-induced STP but not stress-induced grooming or investigative behavior

Next, we probed whether VP signaling may contribute to the differences in grooming behavior or STP following FS in pair-housed versus single-housed female mice. We hypothesized that VP released in the PVN during social interaction may mediate the reduction in grooming and/or STP. One mouse from a female dyad of littermates received an injection of either the V1aR antagonist SR or vehicle 5 min prior to receiving FS (Fig. 2*A*). The stressed subject was then returned to the homecage with the unstressed partner. We assessed behavior as described above. V1aR antagonism had no effect on time spent grooming following stress (Fig. 2*Bi-Cii*). Vehicle- and SR-injected mice spent a similar amount of time self-grooming (vehicle, 307.2 ± 45.9 s, *N* = 5; SR, 241.3 ± 96.1 s, *N* = 5; *p* = 0.55, unpaired *t* test, two-tailed; Fig. 2*D*). This suggests that V1aR signaling is not involved in the social buffering of stress-induced grooming.

**Figure 2.**
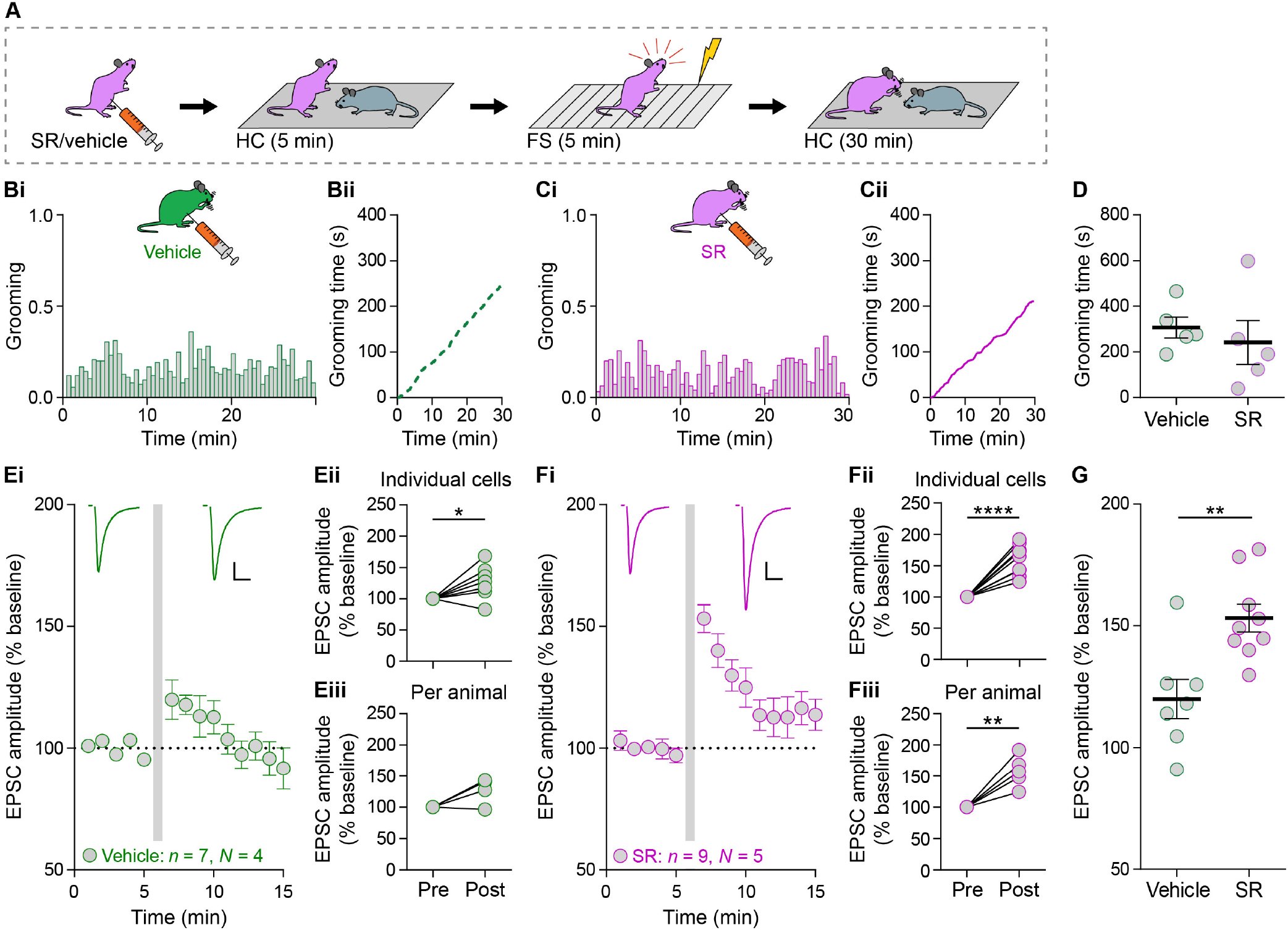
V1aR antagonism prevents social buffering of stress-induced STP but not grooming behavior. **A)** One mouse from a female dyad of littermates was injected i.p. with either the V1aR antagonist SR 49059 (SR, 2 mg/kg) or vehicle 5 min before FS. Immediately following FS, the mouse was returned to its partner in the homecage for 30 min. **Bi)** Histogram showing the amount of self-grooming displayed by vehicle-injected FS mice (*N* = 5) during the 30-min observation period. **Bii)** Cumulative graph illustrates the relative grooming of vehicle-injected FS mice. **Ci)** Histogram showing the amount of self-grooming displayed by SR-injected FS mice (*N* = 5) during the 30-min observation period. **Cii)** Cumulative graph illustrates the relative grooming of SR-injected FS mice. **D)** Vehicle- and SR-injected FS mice spent a similar amount of time selfgrooming (vehicle, mean: 307.2 ± 45.9 s, *N* = 5 mice; SR, mean: 241.3 ± 96.1 s, *N* = 5 mice; *p* = 0.55, unpaired *t* test, two-tailed). Horizontal bars represent the means. **Ei)** EPSCs in CRH^PVN^ neurons of female FS mice injected with vehicle potentiated following HFS (gray bar) relative to baseline. Inset, synaptic currents before and after HFS in vehicle-injected FS mice. **Eii)** STP was present in cells from vehicle-injected FS mice (mean: 119.9 ± 8.1%, *n* = 7 cells, *p* = 0.04 vs. baseline, one-sample *t* test, two-tailed). **Eiii)** STP was absent in vehicle-injected FS mice (mean: 119.7 ± 8.2%, *n* = 4, *p* = 0.09 vs. baseline, one-sample *t* test, two-tailed). **Fi)** EPSCs in CRH^PVN^ neurons of female FS mice injected with or SR potentiated following HFS (gray bar) relative to baseline. Inset, synaptic currents before and after HFS in SR-injected FS mice. **Fii)** STP was present in cells from SR-injected FS mice (mean: 153.2 ± 5.7%, *n* = 9, *p* < 0.0001 vs. baseline, one-sample *t* test, two-tailed). **Fiii)** STP was present in SR-injected FS mice (mean: 154.4 ± 6.4%, *n* = 5, *p* = 0.004 vs. baseline, one-sample *t* test, two-tailed). **G)** STP (individual cells shown) was larger in SR-injected FS mice than in vehicle-injected FS mice (*p* = 0.004, unpaired *t* test, twotailed). Horizontal bars represent the means. Scale bars (**Ei,Fi**) represent 5 ms and 20 pA. Inset currents (**Ei,Fi**) before HFS are scaled to allow for easier comparison after HFS. Error bars represent ± SEM.

Next, we tested the effects of V1aR antagonism at the cellular level. We again performed whole-cell recordings on CRH^PVN^ neurons from vehicle- and SR-injected mice following the 30-min social interaction. STP was observed in CRH^PVN^ neurons recorded from stressed female mice that received vehicle injections (119.9 ± 8.1%, *n* = 7, *p* = 0.04 vs. baseline; Fig. 2*Ei,Eii*) and in neurons from mice injected with SR prior to FS and social interaction (153.2 ± 5.7%, *n* = 9, *p* < 0.0001 vs. baseline; Fig. 2*Fi,Fii*). However, per animal analysis showed that STP was present in SR-injected mice (154.4 ± 6.4%, *N* = 5, *p* = 0.004 vs. baseline; Fig. 2*Fiii*) but was absent in vehicle-injected mice (119.7 ± 8.2%, *N* = 4, *p* = 0.09 vs. baseline; Fig. 2*Eiii*). Furthermore, STP was significantly greater in SR-injected mice than in vehicle-injected mice (*p* = 0.004, unpaired *t* test, two-tailed; Fig. 2*G*). These results indicate the decrease in the synaptic consequences of stress induced by the presence of a partner require V1aRs.

When exposed to a distressed conspecific, rodents engage in unreciprocated investigative behavior directed towards the distressed individual (Knapska et al., 2010; Sterley et al., 2018). One possibility is that this interaction is critical for buffering the synaptic changes in the stressed subject. The total time spent by the unstressed partner spent investigating the stressed subject was unaffected by V1aR blockade (vehicle, 69.3 ± 11.7 s, *N* = 5; SR, 74.9 ± 11.4 s, *N* = 6; *p* = 0.74, unpaired *t* test, two-tailed; Fig. 3*A*). We have previously demonstrated that these investigations target either the head/torso region or the anogenital region of the stressed subject (Sterley et al., 2018). These two behaviors may have different consequences and may differentially contribute to social buffering. We developed a social discrimination index to determine whether V1aRs preferentially affect one feature of the investigative behavior (Fig. 3*C*). Unstressed partners spent a similar amount of time engaged in both anogenital (vehicle, 36.8 ± 6.1 s, *N* = 5; SR, 41.9 ± 6.3 s, *N* = 6; Fig. 3*B*) and head/torso (vehicle, 32.5 ± 5.7 s, *N* = 5; SR, 33.0 ± 5.4 s, *N* = 6; Fig. 3*B*) sniffing toward vehicle- and SR-injected FS mice (one-way ANOVA, F_(3,18)_ = 0.57, *p* = 0.64, followed by Sidak’s multiple comparisons test; Fig. 3*B*). Furthermore, V1aR antagonism did not change the social discrimination index compared to vehicle controls (Fig. 3*C*). Together, these findings suggest that VP signaling is not required for stress-induced grooming in the subject. Further, blocking V1aRs in the subject does not affect investigative behaviors by the partner. The absence of behavioral effects suggest that VP effects on STP may occur locally in the PVN and may target synaptic function very specifically.

**Figure 3.**
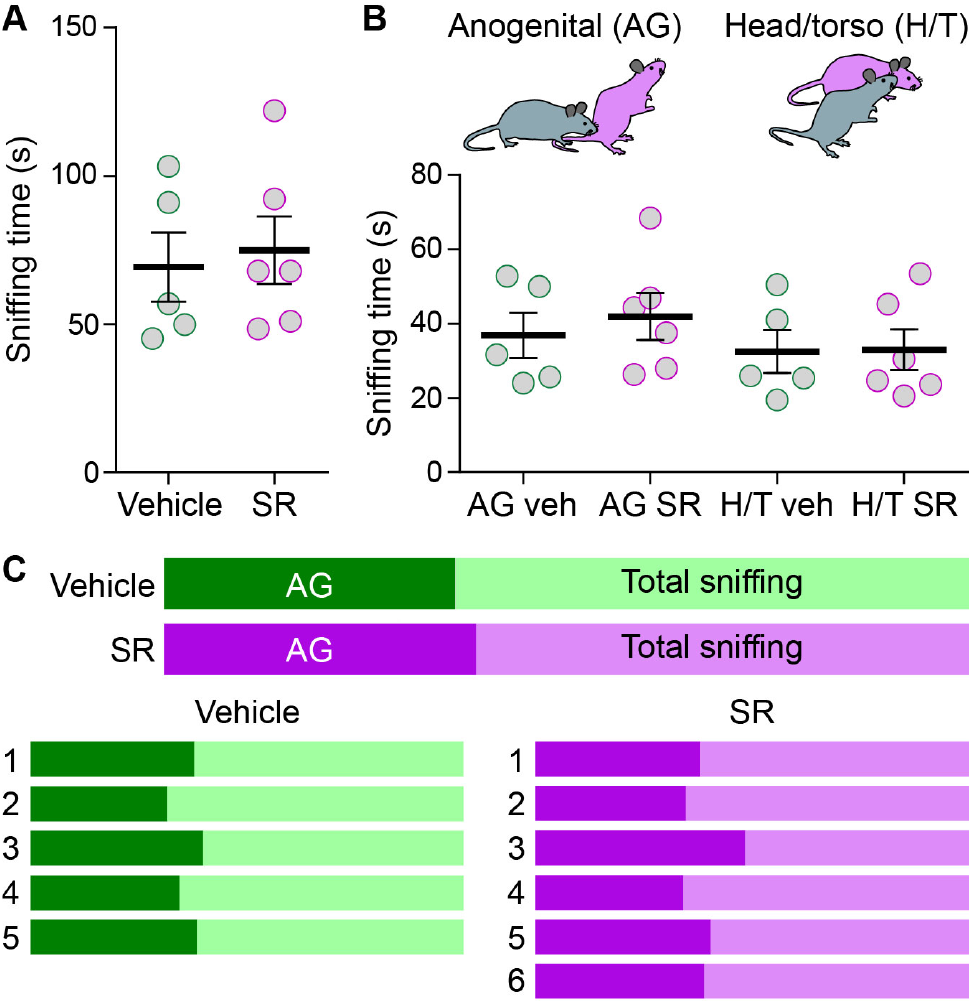
No role for VP receptor signaling in investigative behavior. **A)** Naïve mice spent a similar amount of time investigating vehicle- and SR-injected FS mice (vehicle, mean: 69.3 ± 11.7 s, *N* = 5; SR, mean: 74.9 ± 11.4 s, *N* = 6; *p* = 0.74, unpaired *t* test, two-tailed). **B)** Naïve mice spent a similar amount of time sniffing the anogenital region (AG vehicle, mean: 36.8 ± 6.1 s, *N* = 5; AG SR, mean: 41.9 ± 6.3 s, *N* = 6; *p* = 0.64, one-way ANOVA, F_(3,18)_ = 0.57; *p* = 0.99, followed by Sidak’s multiple comparisons test) and head/torso region (H/T vehicle, mean: 32.5 ± 5.7 s, *N* = 5; H/T SR, mean: 33.0 ± 5.4 s, *N* = 6; *p* > 0.99, Sidak’s multiple comparisons test) of SR- and vehicle-injected FS mice. **C)** Color plots (above) show the overall social discrimination indices for naïve mice toward their respective vehicle- or SR-injected FS partners. Color plots (below) show the social discrimination indices for each individual naïve mouse toward their respective FS partner. Error bars represent ± SEM.

### *Ex vivo* VP reduces stress-induced STP in females but not males

Our *in vivo* experiments above with SR implicate the V1aR in the modulation of stress-induced STP. If inhibition of V1aRs prevents social buffering of STP in CRH^PVN^ neurons, then we would expect VP itself to decrease STP in these cells in the absence of a partner. To test this, we incubated slices from single-housed stressed female mice in VP (100 nM) for 30 min and examined STP. STP was observed in cells recorded from VP-incubated slices (120.2 ± 5.7%, *n* = 16, *p* = 0.002 vs. baseline; Fig. 4*A,B*) and from control aCSF-incubated slices (152.2 ± 8.2%, *n* = 17, *p* < 0.0001 vs. baseline; Fig. 4*A,B*). Per animal analysis gave similar results, with STP being present in VP mice and in aCSF mice (aCSF, 153.8 ± 9.4%, *N* = 14, *p* = 0.0002 vs. baseline; VP, 117.3 ± 5.9%, *N* = 11, *p* = 0.005 vs. baseline; Fig. 4*C*). However, STP in cells from VP-incubated slices was significantly lower than STP in cells from aCSF-incubated slices (*p* = 0.007, one-way ANOVA, F_(2,40)_ = 1.7; Fig. 4*D*). To confirm our *in vivo* findings with the V1aR antagonist, we incubated slices from stressed female mice in SR (10 μM) for 10 min, followed by incubation in both SR and VP for an additional 30 min. Introduction of SR prevented the VP-induced decrease in STP, as STP in SR-incubated slices was significantly higher than STP from VP-incubated slices (*p* = 0.04, Sidak’s multiple comparisons test; Fig. 4*D*) and was similar to that of aCSF-incubated slices (*p* = 0.69, Sidak’s multiple comparisons test; Fig. 4*D*). These findings indicate that activation of V1aRs by VP reduces stress-induced STP in CRH^PVN^ neurons in female mice.

**Figure 4.**
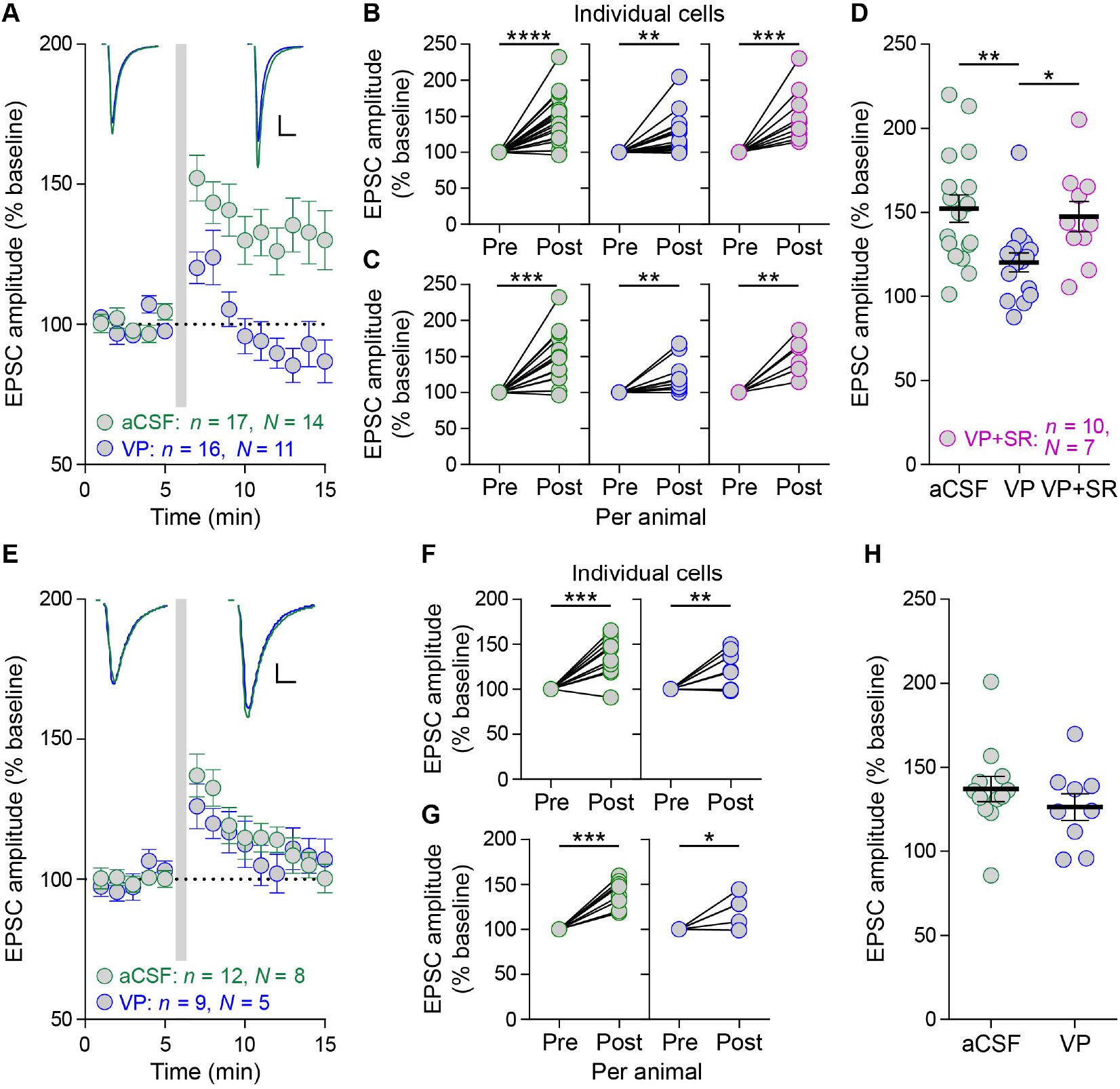
VP reduces STP in CRH^PVN^ neurons from females but not males. **A)** EPSCs in CRH^PVN^ neurons potentiated following HFS (gray bar) relative to baseline in aCSF control slices (green outline) and in VP-incubated slices (blue outline) from single-housed female mice stressed with FS. Inset, synaptic currents before and after HFS in aCSF- and VP-incubated cells. **B)** STP was present in aCSF-incubated cells (mean: 152.2 ± 8.2%, *n* = 17, *p* < 0.0001 vs. baseline, one-sample *t* test, two-tailed), in VP-incubated cells (mean: 120.2 ± 5.7%, *n* = 16, *p* = 0.002 vs. baseline, one-sample *t* test, two-tailed), and in SR+VP-incubated cells (pink outline, mean: 147.5 ± 9.0%, *n* = 10, *p* = 0.0007 vs. baseline, one-sample *t* test, two-tailed). **C)** STP was present female aCSF mice (mean: 153.8 ± 9.4%, *N* = 14, *p* = 0.0002 vs. baseline, one-sample *t* test, two-tailed), in female VP mice (mean: 117.3 ± 5.9%, *N* = 11, *p* = 0.005 vs. baseline, one-sample *t* test, two-tailed), and in female SR+VP mice (mean: 146.8 ± 8.9%, *N* = 7, *p* = 0.001 vs. baseline, one-sample *t* test, twotailed). **D)** STP (individual cells shown) was larger in aCSF-incubated cells (*p* = 0.007, one-way ANOVA, F_(2,40)_ = 1.7; *p* = 0.008, followed by Sidak’s multiple comparisons test) and in SR+VP-incubated cells (*p* = 0.04, Sidak’s multiple comparisons test) than in VP-incubated cells. **E)** EPSCs in CRH^PVN^ neurons potentiated following HFS (gray bar) relative to baseline in aCSF control slices (green outline) and in VP-incubated slices (blue outline) from single-housed male mice stressed with FS. Inset, synaptic currents before and after HFS in aCSF- and VP-incubated cells. **F)** STP was present in aCSF-incubated cells (mean: 137.1 ± 7.6%, *n* = 12, *p* = 0.0001 vs. baseline, one-sample *t* test, two-tailed) and in VP-incubated cells (mean: 126.3 ± 7.9%, *n* = 9, *p* = 0.009 vs. baseline, one-sample *t* test, two-tailed). **G)** STP was present in male aCSF mice (mean: 138.5 ± 5.8%, *N* = 8, *p* = 0.0001 vs. baseline, one-sample *t* test, two-tailed) and in male VP mice (mean: 124.9 ± 9.2%, *N* = 5, *p* = 0.04 vs. baseline, one-sample *t* test, two-tailed). **H)** STP (individual cells shown) in aCSF-incubated cells was similar to that in VP-incubated cells (*p* = 0.35, unpaired *t* test, twotailed). Horizontal bars represent the means. Scale bars (**A,E**) represent 5 ms and 20 pA. Inset currents (**A,E**) before HFS are scaled to allow for easier comparison after HFS. Error bars represent ± SEM.

Our initial observations on social buffering of STP revealed that it is a sex-specific effect that occurs exclusively in female mice (Sterley et al., 2018). As such, we determined whether VP also affects STP in a sex-specific manner by repeating the previous experiments in single-housed male FS mice. STP was observed in CRH^PVN^ neurons from slices incubated in aCSF and in VP (aCSF-incubated, 137.1 ± 7.6%, *n* = 12, *p* = 0.0001 vs. baseline; VP-incubated, 126.3 ± 7.9%, *n* = 9, *p* = 0.009 vs. baseline; Fig. 4*E,F*). Per animal analysis revealed that STP was present in both male aCSF and VP mice (aCSF, 138.5 ± 5.8%, *N* = 8, *p* = 0.0001 vs. baseline; VP, 124.9 ± 9.2%, *N* = 5, *p* = 0.04 vs. baseline; Fig. 4*G*) Finally, there was no difference in the magnitude of STP between aCSF- and VP-incubated cells from stressed male mice (*p* = 0.35, unpaired *t* test, two-tailed; Fig. 4*H*). Collectively, these results suggest that VP selectively buffers stress-induced STP in female mice, similar to the effect of social buffering.

### VP has limited electrophysiological effects on CRH^PVN^ neurons

To further evaluate the effects of VP on CRH^PVN^ neurons, we tested the electrophysiological effects of acute application of VP directly onto CRH^PVN^ neurons from stressed female mice. VP (100 nM) was administered focally to cells via a puff pipette without any blockers in the aCSF. VP had no effect on the firing frequency of action potentials (baseline, 1.0 ± 0.6 Hz; VP, 1.1 ± 0.6 Hz; *n* = 5, *N* = 4; *p* = 0.20; Fig. 5*A,B*) or membrane potential (baseline, –48.3 ± 3.7 mV; VP, –48.6 ± 4.0 mV; *n* = 5, *N* = 4; *p* = 0.63; Fig. 5*A,C*). Next, we analyzed the effects of VP on sEPSCs in CRH^PVN^ neurons following bath application of the peptide in aCSF containing picrotoxin. Similar to the above current clamp experiment, VP (100 nM) had no effect on the frequency (baseline, 7.2 ± 1.8 Hz; VP, 6.5 ± 1.4 Hz; *n* = 5, *N* = 5; *p* = 0.53; Fig. 5*D,E*) or amplitude (baseline, 27.6 ± 4.8 pA; VP, 28.3 ± 5.2 pA; *n* = 5, *N* = 5; *p* = 0.43; Fig. 5*D,F*) of sEPSCs. Taken together, these observations suggest that the electrophysiological effects of VP on CRH^PVN^ neurons following acute stress are specific to synaptic metaplasticity.

**Figure 5.**
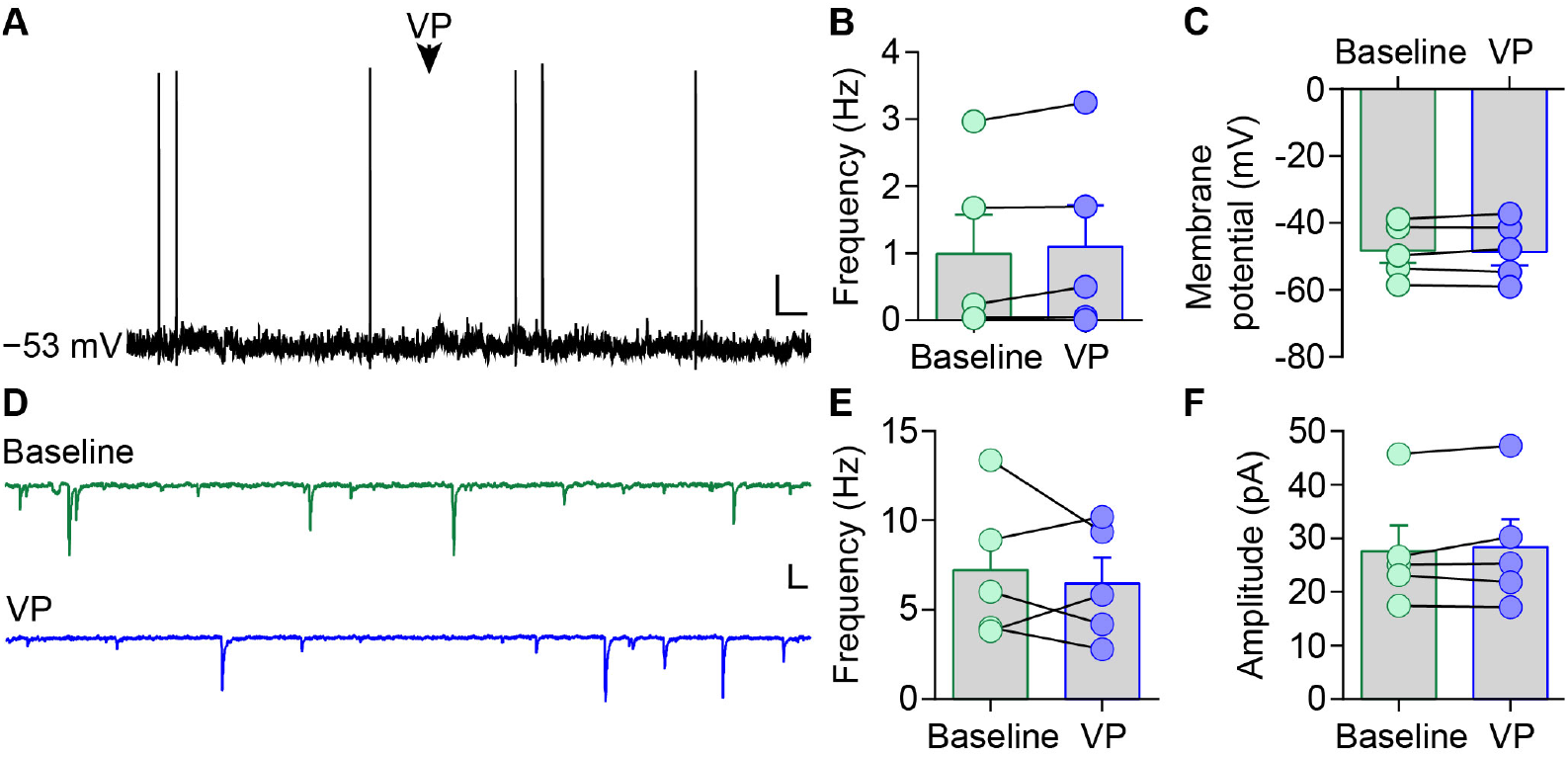
VP has limited electrophysiological effects on CRH^PVN^ neurons following stress. **A)** Representative current-clamp recording from a CRH^PVN^ neuron from a female mouse showing that a VP puff (100 nM, arrowhead) had no effect on the membrane potential or firing. **B, C)** Summary data of the effects of VP on firing frequency (baseline, mean: 1.0 ± 0.6 Hz; VP, mean: 1.1 ± 0.6 Hz; *n* = 5 cells, *N* = 4 mice; *p* = 0.20; **B**) and membrane potential (baseline, mean: −48.3 ± 3.7 mV; VP, mean: *-*48.6 ± 4.0 mV; *n* = 5 cells, *N* = 4 mice; *p* = 0.63; **C**) of CRH^PVN^ neurons from female FS mice. **D)** Representative voltage-clamp traces from a CRH^PVN^ neuron from a female mouse showing that bath application of VP (100 nM) had no effect on the frequency or amplitude of sEPSCs. **E, F)** Summary data of the effects of VP on the frequency (baseline, mean: 7.2 ± 1.8 Hz; VP, mean: 6.5 ± 1.4 Hz; *n* = 5 cells, *N* = 5 mice; *p* = 0.53; **E**) and amplitude (baseline, mean: 27.6 ± 4.8 pA; VP, mean: 28.3 ± 5.2 pA; *n* = 5 cells, *N* = 5 mice; *p* = 0.43; **F**) of sEPSCs in CRH^PVN^ neurons.

### VP acts on CRH^PVN^ neurons to decrease the AMPA/NMDA ratio

Stress-induced metaplasticity is preceded by a decrease in NMDAR-mediated currents in CRH^PVN^ neurons, which can be observed as an increase in the AMPA/NMDA ratio (Kuzmiski et al., 2010). A plausible mechanism for stress buffering by VP is an increase in NMDAR-mediated currents and/or a decrease in AMPAR-mediated currents that restore the AMPA/NMDA ratio toward basal levels. To determine whether VP decreases STP in CRH^PVN^ neurons via modulation of NMDAR- and/or AMPAR-mediated currents, we recorded the AMPA/NMDA ratio in CRH^PVN^ neurons in slices from stressed female mice following VP incubation. The AMPA/NMDA ratio was significantly reduced in cells from VP-incubated slices compared to cells from aCSF control slices (aCSF, 3.8 ± 0.5, *n* = 9; VP-incubated, 2.2 ± 0.4, *n* = 5; *p* = 0.04; Fig. 6*A*). We also recorded the AMPA/NMDA ratio before and after bath application of VP (100 nM) to cells from female FS mice. Acute VP application significantly reduced the AMPA/NMDA ratio within these cells (baseline, 3.3 ± 0.7; VP, 2.4 ± 0.5; *p* = 0.01, *n* = 5, Fig. 6*B*). There was no difference in the AMPA/NMDA ratio between VP incubation and acute VP application (VP-incubated, 2.2 ± 0.4, *n* = 5; acute VP, 2.4 ± 0.5, *n* = 5; *p* = 0.66; Fig. 6*C*).

**Figure 6.**
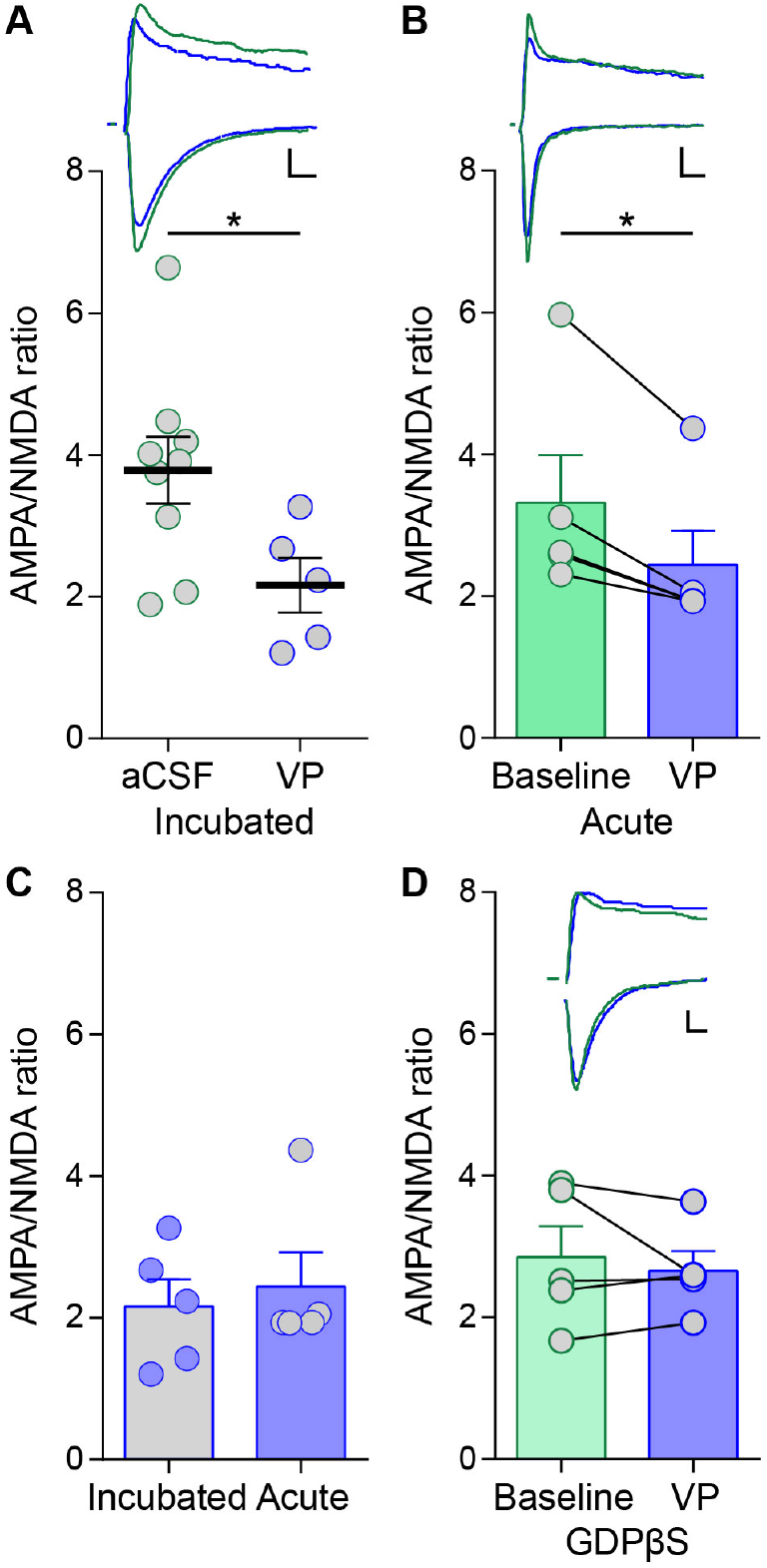
VP decreases the AMPA/NMDA ratio in CRH^PVN^ neurons. **A)** The AMPA/NMDA ratio (individual cells shown) was larger in aCSF-incubated cells (green outline, mean: 3.8 ± 0.5, *n* = 9 cells, *N* = 6 mice) than in VP-incubated cells (blue outline, mean: 2.2 ± 0.4, *n* = 5 cells, *N* = 3 mice; *p* = 0.04, unpaired *t* test, two-tailed) from female mice stressed with FS. Horizontal bars represent the means. Inset, AMPA and NMDA currents from aCSF- and VP-incubated cells. **B)** The AMPA/NMDA ratio was decreased following acute application of 100 nM VP within the same cells (baseline, mean: 3.3 ± 0.7; VP, mean: 2.4 ± 0.5, *n* = 5; *p* = 0.01, paired *t* test, two-tailed). Inset, AMPA and NMDA currents from the same cell before and after VP application. **C)** The AMPA/NMDA ratio was similar between VP-incubated cells and cells following acute VP application (VP-incubated, mean: 2.2 ± 0.4, *n* = 5; acute VP, mean: 2.4 ± 0.5, *n* = 5; *p* = 0.66, unpaired *t* test, two-tailed). **D)** Acute VP application had no effect on the AMPA/NMDA ratio in cells loaded with 1 mM GDPβS (baseline, mean: 2.9 ± 0.4; VP, mean: 2.7 ± 0.3, *n* = 5; *p* = 0.51, paired *t* test, two-tailed). Inset, AMPA and NMDA currents before and after VP application in a cell loaded with GDPβS. Scale bars (**A,B,D**) represent 5 ms and 20 pA. Error bars represent ± SEM.

Our previous experiments with the V1aR antagonist suggest that VP is acting on this receptor to produce its effects on STP. It is not clear whether VP acts directly through V1aRs on CRH^PVN^ neurons themselves, or, as recently shown, indirectly through receptors on astrocytes (Haam et al., 2014). To determine the synaptic location of the V1aRs that mediate the reduction in the AMPA/NMDA ratio, we included GDPβS (1 mM) in the patch pipette to disrupt G protein-mediated signaling in postsynaptic CRH^PVN^ neurons and repeated the experiment described above. Acute VP application had no effect on the AMPA/NMDA ratio in cells loaded with GDPβS (baseline, 2.9 ± 0.4; VP, 2.7 ± 0.3, *n* = 5; *p* = 0.51, Fig. 6*D*). Together, these data suggest that VP acts on V1aRs on CRH^PVN^ neurons, causing a decrease in the AMPA/NMDA ratio and a subsequent reduction in STP.

## Discussion

Our results demonstrate that the presence of a partner buffers the synaptic consequences of stress, and decreases stress-induced grooming behavior in female mice subjected to FS stress. V1aRs are required for social buffering of the synaptic changes but not the grooming behavior. These synaptic actions rely on V1aRs on CRH^PVN^ neurons that reduce AMPA:NMDA and decrease STP. Finally, the local effects of VP on STP are sex-specific, with no local effects on glutamate plasticity in males. These findings establish a critical role for local VP signaling in the PVN in female mice that results in synaptic buffering after stress and social interaction.

Although CRH^PVN^ neurons are traditionally viewed as neuroendocrine cells essential for HPA axis activation, increasing evidence has implicated these neurons in the control of stress-related behaviors (Füzesi et al., 2016; Zhang et al., 2017; Sterley et al., 2018; Daviu et al., 2020). We explored stress-induced grooming in females, an action that rodents initiate following stress (van Erp et al., 1994; Kruk et al., 1998) and one that has been shown to be mediated by collateral projections of CRH^PVN^ neurons to the lateral hypothalamus (Füzesi et al., 2016). We show that the presence of a partner reduces the time spent engaged in self-grooming following stress. Our initial hypothesis, that VP release in the PVN may mediate this social buffering of grooming, was rejected by our observations that V1aR inhibition had no effect on this behavior. This observation is consistent with previous findings that intracerebroventricular (ICV) administration of a selective V1aR antagonist does not reduce novelty-induced grooming in rats (Drago et al., 1997). Furthermore, although V1aR inhibition can suppress grooming induced by central VP administration, antagonists for this receptor are unable to reduce grooming induced by other peptides (Meisenberg and Simmons, 1987), suggesting the inhibitory effect on grooming is specific to that mediated by VP. In addition to the lack of an effect on self-grooming, V1aR inhibition did not influence investigative behavior of the naïve partner toward the stressed individual as assessed by sniffing times and our social discrimination index. This is in agreement with our previous findings that return of the stressed individual to the homecage is likely sufficient to trigger investigative sniffing behavior and that this behavior is independent of CRH^PVN^ neuron activity in that stressed animal (Sterley et al., 2018).

When we compared the activity at glutamatergic synapses on CRH^PVN^ neurons, we observed a reduction in stress-induced STP in pair-housed female mice versus single-housed females. Unlike self-grooming, this social buffering of STP was dependent on V1aR signaling, as inhibition of this receptor produced a robust STP that was similar to single-housed animals. Although CRH^PVN^ neuron activation can drive grooming behavior (Füzesi et al., 2016), our observed increase in STP but lack of an effect on grooming following V1aR inhibition suggests that stress-induced grooming is independent of STP in these neurons. Furthermore, as STP decays within minutes, it is unlikely to be necessary for the maintenance of grooming behavior throughout the observation period. Thus, effects of VP on social buffering are likely localized to the synapse. Although VP is a well-known ACTH secretagogue at the median eminence (Gillies et al., 1982; Antoni, 1993), several studies have demonstrated an inhibitory effect of VP on the HPA axis at the level of the hypothalamus. ICV administration of VP into the third ventricle produces a concentration-dependent decrease in immunoreactive CRH secretion into the hypophysial portal circulation (Plotsky et al., 1984). Social defeat and forced swimming both cause intra-PVN release of VP that is dissociated from peripheral release and likely exerts an inhibitory effect on ACTH secretion (Wotjak et al., 1996, 1998). Additionally, Wotjak and colleagues (1996) observed no significant effect of a V1 antagonist in the PVN on anxiety-related behavior, consistent with our lack of an effect of SR on stress-induced grooming.

While hybridization histochemistry experiments have revealed V1aRs localized to the parvocellular PVN (Ostrowski et al., 1994), previous studies have demonstrated a retrograde neuronal-glial autoregulatory circuit within the PVN mediated by dendritically-released VP (Haam et al., 2014; Chen et al., 2019), implicating astrocytic V1aRs in local VP signaling. Our finding that loading CRH^PVN^ neurons with GDPβS blocks the effects of VP on the AMPA/NMDA ratio suggests that VP mediates its cellular effects through receptors located on CRH^PVN^ neurons themselves. This result also indicates the potential mechanism underlying the VP-mediated reduction in STP. As depression of NMDARs is required for STP (Kuzmiski et al., 2010), our observed decrease in the AMPA/NMDA ratio following VP administration may indicate a restoration of NMDAR function that facilitates the reduction in STP.

We also probed for the effects of VP on STP in males. Unlike our results from female mice, we observed no change in STP following VP incubation of slices from male animals. Although we did not explore the mechanism underlying these sex differences, it is unlikely that differences in VP release between males and females account for this sexually dimorphic effect since VP incubation did not change STP in males. Instead, these experiments suggest a difference that is downstream of VP release, but does not completely negate the possibility of a concurrent upstream difference. One possible explanation relates to V1aR desensitization. Like many GPCRs, the V1aR undergoes agonist-induced phosphorylation by G protein-coupled receptor kinases, which promotes β-arrestin association and a subsequent internalization of the receptor (Innamorati et al., 1998, 1999; Bowen-Pidgeon et al., 2001). While no studies have reported sex differences in this process for V1aRs, an example of this does exist for CRH receptor 1 (CRHR1) in locus coeruleus neurons. In male rats, acute swim stress causes β-arrestin 2 to associate with CRHR1 and leads to internalization of the receptor; however, in females, this stress-induced internalization is impaired due to diminished CRHR1-β-arrestin 2 association, which renders these neurons more sensitive to CRH (Bangasser et al., 2010). It is possible that a similar sex difference exists for V1aRs in CRH^PVN^ neurons.

Sex differences in VP signaling have also been attributed to polymorphisms in repetitive DNA sequences, or microsatellites, in the 5’ regulatory region of the gene encoding the V1aR, *Avpr1a.* Of particular interest are two highly polymorphic microsatellites in the promotor region, RS1 and RS3. Microsatellite length predicts intraspecific variation in V1aR expression and social behavior in male prairie voles, with long-allele males displaying increased social investigation and bonding behavior compared to short-allele males (Hammock and Young, 2005). Both short- and long-allele males show robust differences in V1aR binding in several brain regions (Hammock and Young, 2005). In humans, longer RS3 repeat length is associated with increased altruistic behavior and higher V1aR mRNA levels in the hippocampus (Knafo et al., 2008). Polymorphisms in *Avpr1a* in the PVN that lead to changes in receptor expression may account for our observed sex differences in the effects of VP on STP in CRH^PVN^ neurons. Future experiments using genetic analysis are required to confirm the existence of these polymorphisms in the PVN of mice.

The fundamental question that remains is what causes VP release in the PVN during social buffering. One intriguing mechanism is through the odorant receptor 37 (OR37) subfamily, part of an olfactory subsystem that has been previously implicated in stress buffering. Endogenous ligands for OR37 receptors, which are found in bodily secretions from mice, reduce c-Fos expression in CRH^PVN^ neurons following exposure to a novel environment (Klein et al., 2015). In humans, exposure to undetected hexadecanal, a ligand for OR37 receptors, reduces the acoustic startle response in typically-developed adults but has no effect in cognitively-able adult participants with autism spectrum disorder (ASD) (Endevelt-Shapira et al., 2018). The mitral cells of OR37 glomeruli project specifically to VPergic cells in the PVN (Bader et al., 2012). As both magnocellular and parvocellular VP neurons are capable of dendritic release of VP (Ludwig, 1998), activation of the OR37 subsystem may be the source that initiates VP release in the PVN. However, future experiments are required to confirm an involvement of this subsystem in VP-mediated social buffering.

Our findings have translational implications for several stress-related psychiatric disorders, such as post-traumatic stress disorder (PTSD), anxiety, and depression. Dysfunction of the VP system is often associated with these disorders (Keck, 2006). For instance, urinary VP levels have been both positively (de Kloet et al., 2008) and negatively (Marshall, 2013) correlated with PTSD symptom severity in men. Meanwhile, plasma VP levels are elevated in patients with major depression (van Londen et al., 1997). In addition to stress-related disorders, altered VP signaling is also associated with ASD, of which abnormal social behavior is a main characteristic.

In male children with ASD, lower VP concentration in the CSF predicts greater social impairments (Oztan et al., 2018). In a recent phase 2 clinical trial, intranasal VP administration improved the primary outcome measure of social abilities and diminished anxiety symptoms in children with ASD (Parker et al., 2019). Understanding the cellular substrates and mechanisms that underlie social buffering will help advance the treatment of these disorders.

## Acknowledgments

We thank Cheryl Breiteneder for technical assistance. This work was supported by an operating grant to J.S.B. from the Canadian Institutes of Health Research (CIHR, FDN-148440). S.P.L. is the recipient of a CIHR Doctoral Research Award.

## Notes

**Conflict of interest:** The authors declare no competing financial interests.

### Competing Interest Statement

The authors have declared no competing interest.

